# Putative conjugative plasmids with *tcdB* and *cdtAB* genes in clinical *Clostridium difficile* strains from MLST clades C-I, 2 and 4

**DOI:** 10.1101/852418

**Authors:** Gabriel Ramírez-Vargas, César Rodríguez

**Affiliations:** Facultad de Microbiología, CIET, LIBA, Universidad de Costa Rica, Costa Rica

**Keywords:** *C. difficile*, MGEs, extrachromosomal elements, TcdB, CDT

## Abstract

The major toxins of *Clostridium difficile* (TcdA, TcdB, CDT) are encoded chromosomally in nearly all known strains. Following up on a previous report, we found five new examples of a family of putative conjugative plasmids with *tcdB* and *cdtAB* in clinical *C. difficile* isolates from MLST Clades C-I, 2, and 4.

## Text

Upon gut microbial dysbiosis, ingested *Clostridium difficile* spores may differentiate in the human colon into vegetative cells and release one or two large clostridial cytotoxins (i.e. TcdA, TcdB) and/or a binary toxin with ADP-ribosyltransferase activity (CDT) to cause colitis and diarrhea. This pathogen has gained notoriety in the last decade and is now considered a growing public health threat and burden on healthcare systems (*1*).

Not all *C. difficile* strains are toxigenic, but when present, genes for TcdA, TcdB and CDT are nearly without exception encoded by two separate chromosomal loci known as PaLoc and CdtLoc (*2*). This paradigm has been challenged by the rather recent finding of Clade C-I strains SA10-050, CD10-165 in France (*3*) and HSJD-312, HMX-152 in Costa Rica (*4*), which carry a monotoxin *tcdB*^+^ PaLoc next to a full Cdtloc on extrachromosomal molecules that resemble conjugative plasmids (*4*). In this regard, Clade C-I strains have been traditionally considered to be of environmental origin (*5*) and non-toxigenic due to lack of perfect PaLoc integration sites in their chromosomes (*6*).

The Research Laboratory for Anaerobic Bacteriology (LIBA) has been isolating and typing *C. difficile* in Costa Rica for nearly a decade and thereby generated an isolate collection with over 800 records. A search of mobile genetic elements (MGEs) among Illumina whole genome sequences from 150 of those bacteria, led to the discovery of five new *tcdA*^−^/*tcdB*^+^/*cdtAB*^+^ extrachromosomal DNA molecules among isolates that were recovered between 2013 and 2018 from patients that developed diarrhea at three Costa Rican hospitals (LIBA-6656, LIBA-7194, LIBA-7602, LIBA-7678, LIBA-7697). Raw sequencing data for isolate LIBA-6656 can be retrieved from the European Nucleotide Archive (Run ERR467623). In turn, reads for the other four isolates are available at the MicrobesNG platform (https://microbesng.com/portal/projects/FB43968C-E9EF-4270-9D1A-054457CC9B54/).

As indicated by a tree of aligned, concatenated, *C. difficile* MLST allele combinations deposited in pubMLST (Figure 1a), these putative plasmid sequences were found not only in Clade C-I isolates (LIBA-7194, LIBA-7602, LIBA-7678), but also in isolates from clades 2 (LIBA-6656) and 4 (LIBA-7697). This result expands the host range previously reported by us for this type of MGEs to *C. difficile* clades of more common association with humans, raising stimulating questions about their role in human disease.

**Figure 1.**
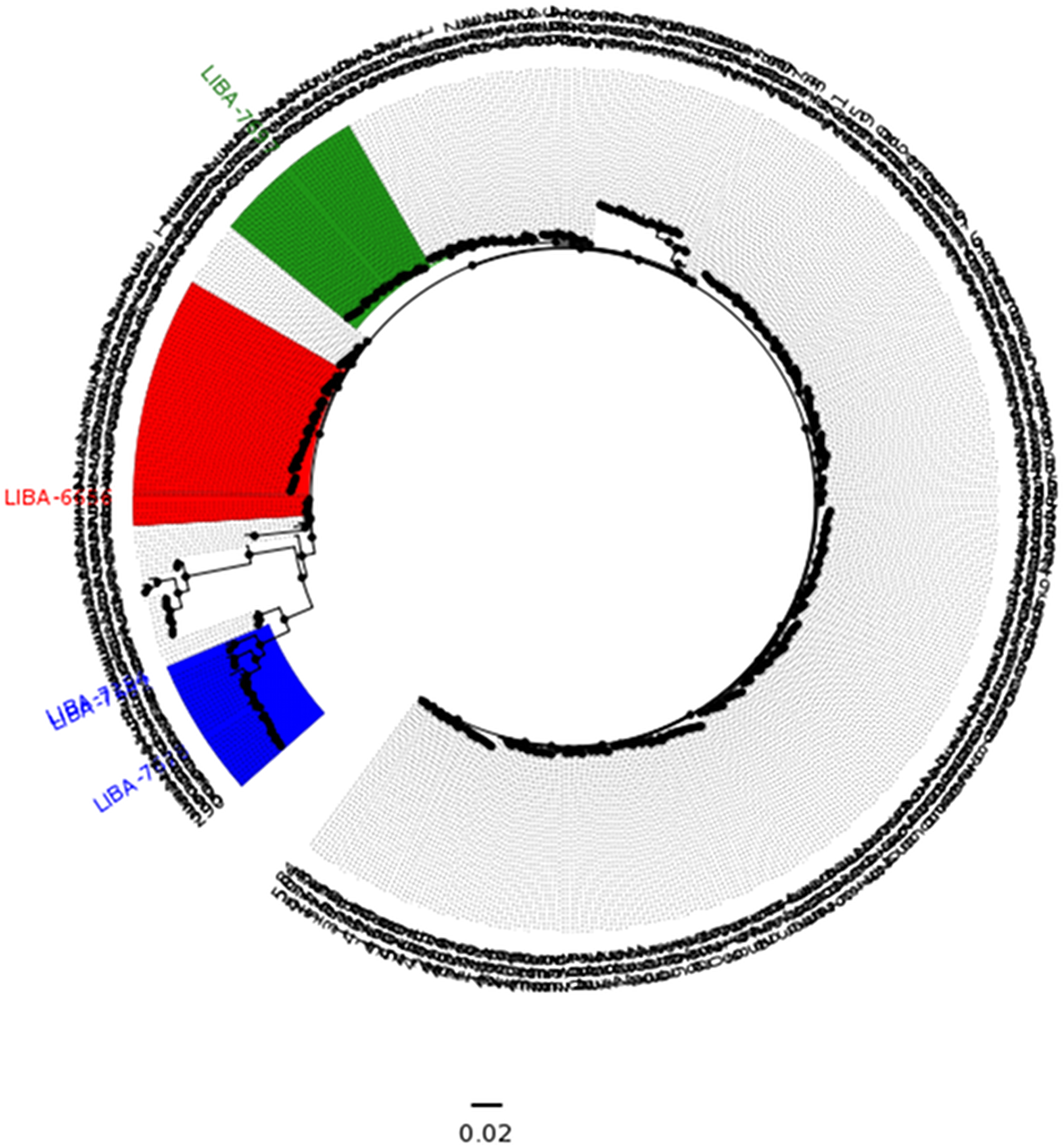
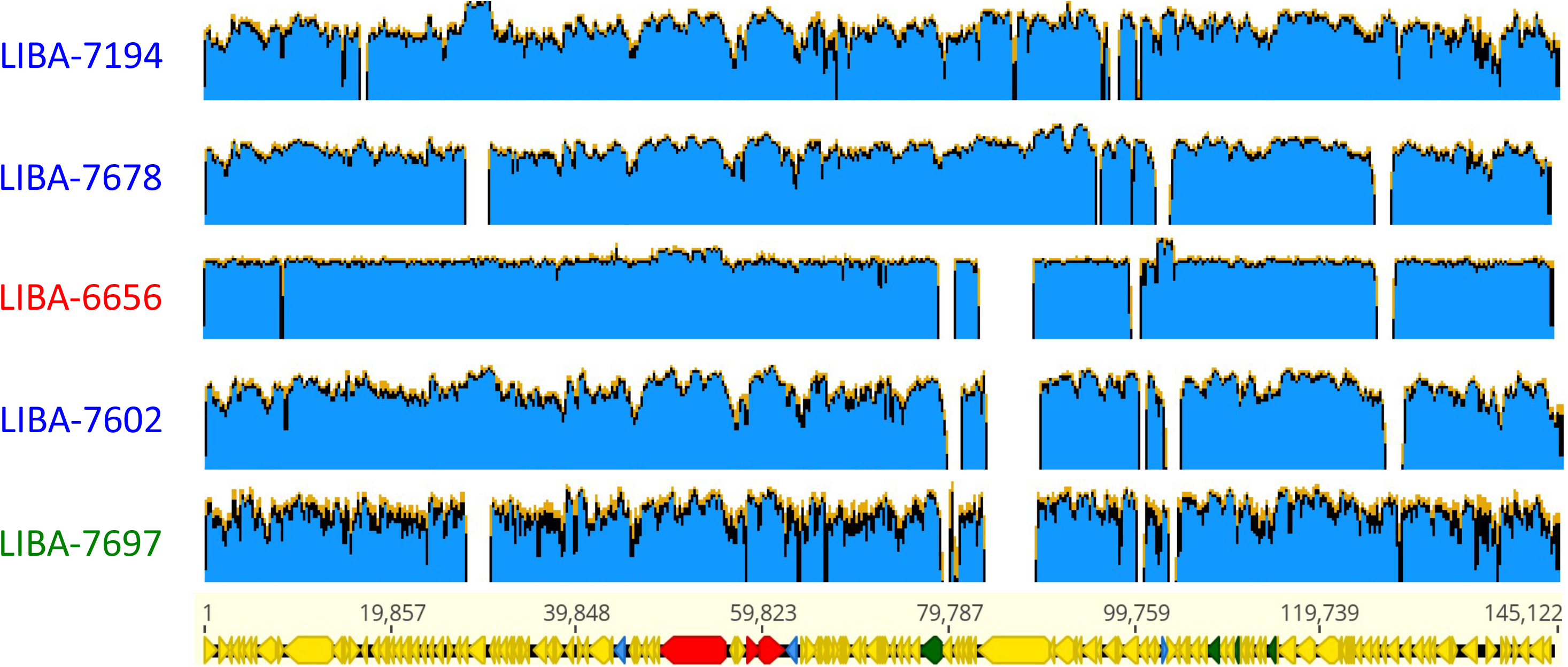
MLST-based classification (A) and diversity of extrachromosomal circular sequences (B) of *C. difficile* strains with plasmid-encoded toxins. The FastTree tree shown in (A) was derived from a MUSCLE alignment of concatenated MLST alleles from all *C. difficile* sequence types (STs) deposited in the PubMLST database. Tip labels represent STs or strain names. Strains from clades C-I, 2, and 4 are highlighted in blue, red, and green, respectively. Panel B shows short reads from strains LIBA-7194, LIBA-7678, LIBA-7602 (Clade C-I, blue), LIBA-6656 (Clade 2, red), and LIBA-7697 (Clade 4, green) mapped to the plasmid sequence of strain HSJD-312, which was obtained through long-read PacBio sequencing and therefore used as a reference (145 kb, bottom). Arrows in the reference sequence represent annotated CDS. Different colors were used to show genes for toxins (red), transposases, integrases, and recombinases (blue), and proteins from a putative conjugation machinery (green).

BWA mapping of reads from isolates LIBA-6656, LIBA-7194, LIBA-7602, LIBA-7678 and LIBA-7697 to a plasmid sequence obtained by PacBio sequencing of strain HSJD-312 (Figure 1b) revealed that the new plasmid sequences are related, yet not identical to each other (92-98% coverage) or to the circular extrachromosomal sequences previously obtained for Clade C-I strains HSJD-312 and HMX-152 (91-98% coverage). Whereas their toxin loci, agr locus, and potential conjugation machinery were conserved, mapping gaps corresponded to putative virulence factors (i.e. putative lectin-or cell wallbinding proteins), hypothetical proteins, and MGEs, such as a class 2 intron and transposases (Figure 1b). This data matches the anticipated mobility potential of the *C. difficile* toxin plasmids (*4*) and supports the notion that they belong to a family of chimeric molecules undergoing non-homologous recombination (*4*).

The MGE-associated *tcdB* sequence of the clade 2 strain LIBA-6656 could not be fully assembled. In the remaining four strains this gene was highly conserved and expected to encode variant TcdBs (99-100% protein sequence identity) that would cause a “sordellii” cytophatic effect (*7*). Besides its plasmidial *tcdB*, LIBA-6656 carries a different *tcdB* on a chromosomal PaLoc. This surprising finding demonstrates that two *tcdB* alleles can coexist in a single strain. The contribution of each *tcdB* to infection is unclear at this moment, yet the coexistence of two PaLocs within a host is compatible with the suggested transition from ancient monotoxin PaLocs to modern bitoxin PaLocs (*3*). A high level of sequence identity was also noted for *cdtA* (≥99%) and *cdtB* (≥98%) in all five putative plasmids. As previously reported, the toxin genes of the new putative plasmid sequences are flanked by recombinases and integrases (*4*). Other elements from this family of potential MGEs lack toxin genes (4), indicating that the latter were likely gained through lateral gene transfer events. However, it is difficult with such a small dataset to determine whether the noted conservation of toxin gene sequences reflects stable coevolution or simply short evolutionary time after their acquisition.

Three of the five isolates that host new toxin plasmids would have remained undetected if we had not attempted *C. difficile* cultivation from TcdB^−^ stool samples of patients with CDI symptoms and sequenced isolates with negative results for *tcdC* and *tcdA* in a PCR-based screening (LIBA-7194, LIBA-7602, LIBA-7678). Hence, we anticipate that the frequency of *C. difficile* isolates with toxin plasmids has been underestimated and urge for refined diagnostic procedures. Moreover, our results open avenues to explore whether plasmids from this group are present in species other than *C. difficile* and account for undiagnosed cases of antibiotic-associated diarrhea.

## Acknowledgements

Vicerrectoría de Investigación/UCR funded this work. Dr. Thomas Riedel and Prof. Jörg Overmann generated the PacBio plasmid sequence of isolate HSJD-312 that was used as a reference in the context of a BMBF-MICITT project.

